# A mutation-selection model of protein evolution under persistent positive selection

**DOI:** 10.1101/2021.05.18.444611

**Authors:** Asif U. Tamuri, Mario dos Reis

## Abstract

We use first principles of population genetics to model the evolution of proteins under persistent positive selection (PPS). PPS may occur when organisms are subjected to persistent environmental change, during adaptive radiations, or in host-pathogen interactions. Our mutation-selection model indicates protein evolution under PPS is an irreversible Markov process, and thus proteins under PPS show a strongly asymmetrical distribution of selection coefficients among amino acid substitutions. Our model shows the criteria *ω* > 1 (where *ω* is the ratio of non-synonymous over synonymous codon substitution rates) to detect positive selection is conservative and indeed arbitrary, because in real proteins many mutations are highly deleterious and are removed by selection even at positively-selected sites. We use a penalized-likelihood implementation of our model to successfully detect PPS in plant RuBisCO and influenza HA proteins. By directly estimating selection coefficients at protein sites, our inference procedure bypasses the need for using *ω* as a surrogate measure of selection and improves our ability to detect molecular adaptation in proteins.

**Significance Statement:** Understanding how natural selection acts on proteins is important as it can inform studies from adaptive radiations to host-pathogen co-evolution. Traditionally, selection on proteins is inferred indirectly by measuring the non-synonymous to synonymous rate ratio, *ω*, with *ω* > 1, = 1, and < 1 indicating positive (adaptive) selection, neutral evolution, and negative (purifying) selection respectively. However, the theoretical underpinnings of this ratio are not well understood. Here we use first-principles of population genetics to work out how persistent changes in selection affect protein evolution and we use our new model to detect adaptive sites in plant and influenza proteins. We show measuring selection directly improves detection of adaptation in proteins.

---

**U**nderstanding how natural selection acts on molecular sequences has long been a pursuit of evolutionary biology. For example, Kimura (1), using a model that assumes the genome has an infinite number of sites, showed the relative rate of molecular evolution is approximately given by

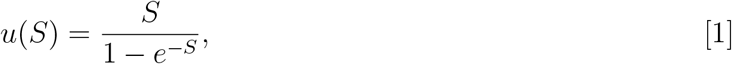

where *S* is the selection coefficient acting on mutations. If new mutations in the genome are positively selected (*S* > 0) the relative rate of molecular evolution is accelerated (*u* > 1), while the rate is the neutral mutation rate (*u* = 1) if there is no selection (*S* = 0), and the rate is decelerated (*u* < 1) if mutations are negatively selected (*S* < 0).

Equation 1, which is the relative probability of fixation of selected over neutral mutations (2–5), has important implications for understanding molecular adaptation in proteins. For a sample of protein-coding sequences from various species, the ratio between the number of substitutions at non-synonymous sites (which are under selection) and at synonymous sites (which are under weak or no selection) should approximately follow the dynamics of Eq. 1 (6). This ratio, commonly known as *ω* = *d_N_ / d_S_*, is widely used as a test of molecular adaptation in proteins, with *ω* >, *ω* = 1 and *ω* < 1 interpreted as evidence of molecular adaptation (positive selection), neutral evolution, and purifying selection respectively.

However, Kimura’s relative rate of molecular evolution (Eq. 1), based on the infinite-sites model (7, 8), assumes all new mutations appear at new sites in the genome. This assumption appears unrealistic for proteins. Nielsen and Yang (6) have argued that if amino acid fitnesses are re-assigned every time a new mutation appears at a site in a protein (so that the selection coefficient, *S*, is always the same at the site), then Eq. 1 gives the relationship of *S* and *ω* under a finite-sites model. However, it is not clear in which condition this fitness reassignment should apply: If an *i* to *j* mutation has selection coefficient *S*, then the reverse *j* to *i* mutation should have coefficient − *S*, but Nielsen and Yang’s model assumes it reverts to *S*. Without this assumption it does not appear possible to equate *ω* = *u(S)*.

Spielmann and Wilke (9), and dos Reis (10), used the Fisher-Wright mutation-selection model (2, 3, 11) to derive the relationship between *ω* and the selection coefficients acting on codon sites within a finite-sites model. They showed that *ω* ≤ 1 when selection coefficients are constant over time (i.e. they are not reassigned, (9, 10)); and *ω* > 1 for a short period of time after selection coefficients undergo a single shift during an adaptive event, for example, when a virus adapts to a new host (10).

However, the relationship between *ω* and selection coefficients under the more general case of persistent changes in selection over time appears unclear. This case, which we term persistent positive selection (PPS), is important because selection coefficients acting at codon sites may change repeatedly during persistent environmental changes, during adaptive radiations, and in host-pathogen interactions (such as in a virus evading herd immunity in a host population). Thus, understanding how PPS affects *ω* in proteins can inform the development of methods to detect positive selection and give us insight onto the mechanisms of adaptive evolution in general.

Here we develop a mutation-selection model of codon substitution under PPS. Analysis under the new model indicates codon substitution is an irreversible Markov process, leading to a highly asymmetrical distribution of selection coefficients among substitutions in proteins under PPS. More strikingly, the PPS model shows the criteria *ω* > 1 to detect molecular adaptation in proteins is conservative and indeed arbitrary. We demonstrate, using empirical data, that PPS sites in proteins may have substitution rates that are lower than the neutral rate (i.e., *w* < 1).

## Theory

### The PPS codon substitution model

We develop the new model by integrating the non-homogeneous selection model of Kimura and Ohta (12) with the mutation-selection codon substitution model of Halpern and Bruno (11). Consider a population of organisms with haploid genome number *N*. That is, the number of copies of the genome in the population is *N* (i.e., the population size is *N* if the organism is haploid and *N/*2 if it is diploid). Suppose a site *k* in a protein-coding gene is fixed for codon *i* in the population, and the scaled Malthusian fitness of *i* is *F_i,k_*. A new mutant codon *j* appears at the site and has initial selective advantage 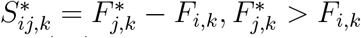. The selective advantage then decays exponentially as a function of time (12), for example, due to gradual environmental change. Kimura and Ohta (12) showed that the fixation probability of *j* is approximately 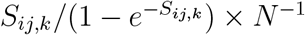 where *S_ij,k_* is constant and 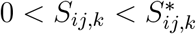. In other words, the fixation probability of *j* is the same as that of an allele with intermediate, but constant, selective advantage *S_ij,k_*.

It appears that other types of decay function lead to the same fixation probability. For example, the same result is obtained in the case of frequency-dependent selection (FDS) when the fitness of *j* decays exponentially as a function of the frequency of *j* in the population (13). In the case of FDS, once *j* becomes fixed, any new mutant alleles may have high fitnesses because they would be rare. We expect this type of dynamics in, for example, viruses escaping the herd immunity of a host population. Similarly, if the environment gradually shifts between two states, then the selective advantage of *j* or *i* would be continuously reset depending on the particular environment. This would then lead to re-setting (or re-assignment) of the fitnesses of *i* and *j*. This persistent change in the selection coefficient is what we term persistent positive selection (PPS). We formalize the PPS dynamics next.

Let the selection coefficient for the *i → j* mutation be *S_ij,k_ = F_j,k_ − F_i,k_ + Z_k_*, where *F_j,k_*, *F_i,k_* and *Z_k_*(≥ 0) are constant. Let the selection coefficient for the reverse mutant, *j → i*, be *S_ji,k_ = F_i,k_ − F_j,k_ + Z_k_*. In other words, we have partitioned the fitnesses of *j* and *i* into two components: A constant component, *F_j,k_* and *F_i,k_*, representing structural constrains of the protein on the amino acid encoded by the codon; and *Z_k_*, the PPS component. Thus, when *Z_k_* > 0, the selection coefficient is persistently reset with new mutations.

The substitution rate from *i* to *j* at location *k*, *q_ij,k_*, is equal to the neutral mutation rate, *μ_ij_*, times the number of *i* alleles in the population, *N*, times the fixation probability of the *j* mutant (1, 11). Assumming synonymous substitutions are neutral, this gives

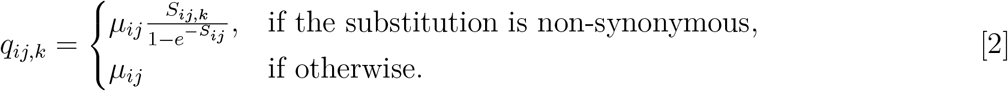

### Irreversibility of codon substitution under PPS

Eq. 2 describes codon substitution as a continuous Markov process. Polymorphisms are ignored and the population is assumed to switch from *i* to *j* instantaneously. This assumption appears reasonable if 2*Nμ_ij_* ≪ 1, for all *μ_i,j_*. The proportion of time location *k* remains fixed for *j* (i.e. the stationary frequency of *j*) is *π_j,k_*. A Markov process is said to be reversible in equilibrium if it satisfies the detailed-balance condition *π_i,k_q_ij,k_ = π_j,k_q_ji,k_* (14). When *Z_k_* = 0, the model of Eq. 2 is reversible (15). However, when *Z_k_* > 0 the process is, in general, irreversible because the detailed balance condition does not hold. When *Z_k_* > 0, the stationary frequencies are found by solving the system of equations Σ *π_j,k_q_j,i,K_* − Σ *π_j,k_q_j,i,K_* with the constraint Σ *π_i,k_* = 1. We calculate the irreversibility index for site *k* as *I_k_ = |π_i,k_q_ij,k_ − π_j,k_q_ji,k_*|, where *I_k_* > 0 indicates evolution at site *k* is irreversible, and *I_k_* = 0 otherwise (16).

### Identifying protein locations under PPS

Given an alignment of protein-coding genes with corresponding phylogeny, the model of Eq. 2 can be used to estimate the fitnesses of amino acids at each site using maximum penalised likelihood, as we have done previously with the simpler version of the model with *Z_k_* = 0 (17). For each site in the alignment, we compare the null model *Z_k_* = 0 (no PPS) against *Z_k_* > 0 (PPS) using a likelihood-ratio test. Because of the boundary condition (*Z_k_* > 0) in the test and the use of penalised likelihood, the distribution of the likelihood-ratio statistic does not follow the typical *χ*^2^ distribution. Thus, we use Cox (18) simulation approach as used in phylogenetics (19) to obtain the appropriate null distribution (see Methods). Our simulation analysis indicates the method has excellent behaviour to detect PPS in simulated data (Table 1). For example, for *Z* >= 5, the method can detect over 94% of all PPS sites and its robust to the penalty parameter used to estimate *Z* (Table 1).

**Table 1.** Performance of the LRT for detecting PPS sites in simulated data after FDR correction (5%)

| True model | $\lambda = 0.01$ | $\lambda = 0.5$ | $\lambda = 1.0$ |
| --- | --- | --- | --- |
| **FPR at 0.05 significance** |  |  |  |
| swMutSel ($Z = 0$) | 0.066 | 0.066 | 0.066 |
| **TPR at 0.05 significance** |  |  |  |
| swMutSel+PPS ($Z = 2$) | 0.441 | 0.452 | 0.449 |
| ($Z = 5$) | 0.952 | 0.952 | 0.947 |
| ($Z = 10$) | 0.965 | 0.963 | 0.960 |

FPR: False positive rate, TPR: True positive rate.

**Table 2.** Number of sites estimated to be under PPS in three real datasets.

| Dataset | # taxa | # sites | # $Z > 0$ ($\omega > 1$) |
| --- | --- | --- | --- |
| Plant RBCL | 478 | 466 | 65 (55) |
| Influenza HA | 466 | 589 | 18 (18) |
| Mammal CYTB | 418 | 407 | 0 (0) |

### The relationship between selection coefficients and *ω*

The average substitution rate of codon site *k*, averaged over time is

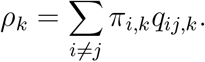

This rate can be separated into its non-synonymous and synonymous components, *ρ_k_ = ρ_N,k_ + ρ_S,k_*, where

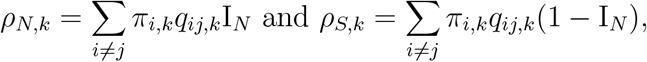

and where the indicator function I*_N_* = 1 if the *i* to *j* substitution is non-synonymous, and = 0 if otherwise. For a neutrally evolving sequence (e.g. apseudogene) the corresponding rates are

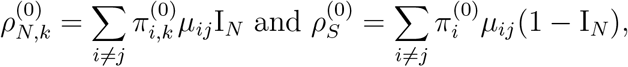

where 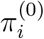 is the stationary frequency of *i* without selection. Thus, the relative non-synonymous rate is

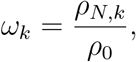

((10), see (9), for a slightly different definition of *ω_k_*).

## Detection of PPS in real proteins

We tested for PPS sites in three real sequence datasets: the haemagglutinin protein (HA) from human influenza H1N1 virus, the rbcL protein subunit from flowering plants, and the mitochondrial cytochrome b (CYTB) protein from mammals (Table 1). Given the multiple sequence alignment, phylogeny and mutational parameters we estimated the sitewise amino acid fitnesses *F* and the PPS component *Z*. We then performed the LRT of PPS vs no PPS and used false discovery rate (FDR) at the 5% level to identify sites under PPS. We detected PPS (*Z* > 0) at 65 sites in the plant RBCL and 18 sites in the influenza HA, but we find no PPS sites in mammal CYTB (Table 1). Interestingly, only 55 out 65 of PPS sites in RBCL have *ω* > 1. For HA, all 18 PPS sites also have *ω* > 1. The location of PPS sites and estimated *ω_k_* values are shown in Fig. 1A–A”.

**Fig. 1.**
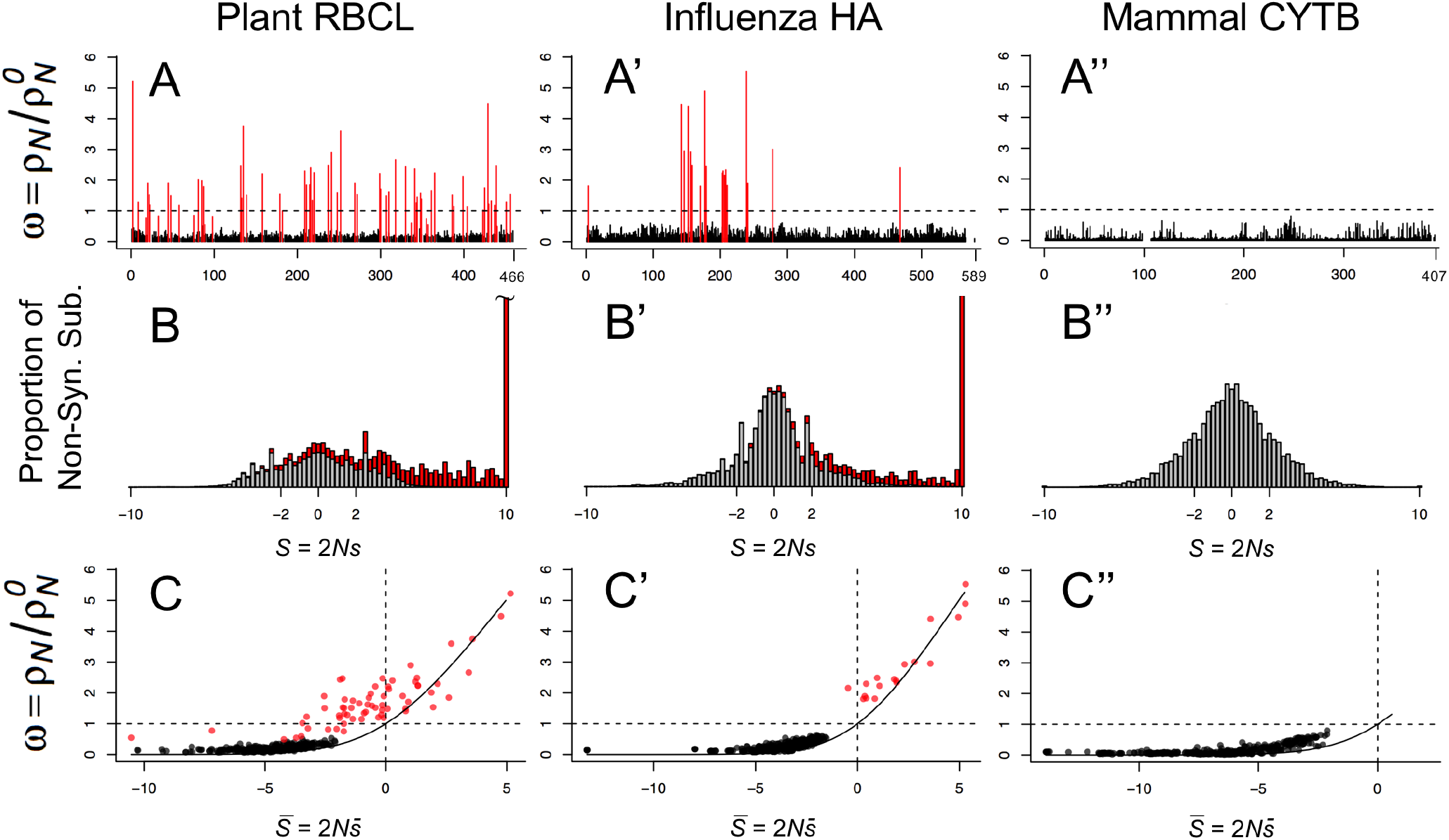
Analysis of proteins under the PPS mutation-selection model. (A-A”) Estimates of *ω* at protein sites. (B-B”) Distribution of selection coefficients among non-synonymous substitutions. (C-C”) Relationship between *ω* and average selection 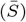 at protein sites. Sites under PPS (*Z_k_* > 0) are indicated in red in A-A” and C-C”, and their contribution to the distribution of selection coefficients indicated in red in B-B”. In C-C’ the solid line is Eq. 1

### The distribution of selection coefficients at sites under PPS is asymmetrical

We estimated the distribution of selection coefficients among non-synonymous substitutions (17) in the three protein-coding genes analysed (Fig. 1B–B“). For non-PPS sites (i.e. sites where *Z* = 0), the distribution of selection coefficients is symmetrical, with a mode at *S* = 0, because under the detailed balance condition the proportion of slightly deleterious mutations fixed in the population over time is equal to the proportion of advantageous mutations fixed (15). However, among PPS sites in plant RBCL and influenza HA, the distribution is highly skewed with a mode at *S* > 10 because for these sites, irreversibility of the substitution process means the detailed balance condition does not apply, and thus there is a persistent excess of advantageous mutations being substituted into the population. The asymmetry in the distribution is more marked for sites with *Z_k_* >> 0 as those sites tend to show the strongest deviation from detailed balance (Fig. 2).

**Fig. 2.**
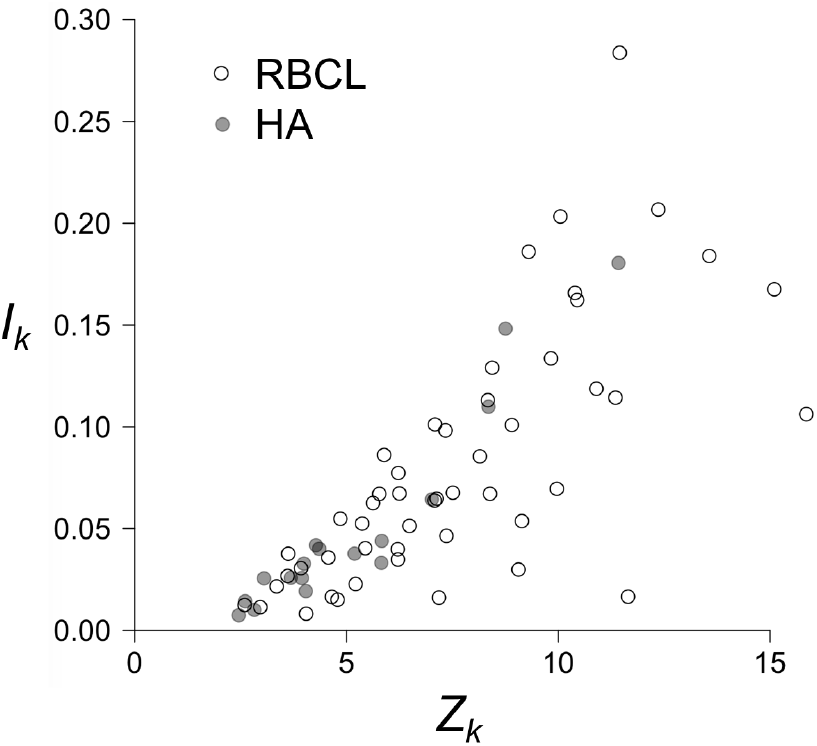
Irreversibility index, *I_k_*, v *Z_k_* for protein sites under PPS. *I_K_* measures the deviation of the substitution process from detailed balance, with *I_K_* = 0 indicating a reversible process and *I_k_* > 0 an irreversible one.

### PPS sites are under strong purifying constraints

At equilibrium, the average selection coefficient of new mutations at site *k* is

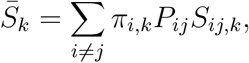

where *P_ij_ = μ_ij_/ Σ_j_ μ_ij_* is the probability that the next mutation is *j* given the site is currently fixed for *i* (10). If most new mutations are very deleterious, then the site is under purifying selection and 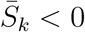, while if most new mutations are advantageous the site is under diversifying selection and 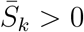. Historically, *ω_k_* has been used as a proxy for 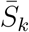, based on the approximation of Eq. 1 (20). Thus calculating 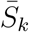 should provide insight into the relationship between the strength of selection at a site and *ω_k_*.

Fig. 1C–C′ shows the estimated 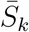 for the three datasets plotted against *ω_k_*. For 43 PPS sites in RBCL and one PPS site in HA, we find that 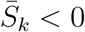. This shows PPS sites are effectively under a mixture of purifying selection against deleterious amino acid substitutions, and diversifying selection in favor of a few amino acids that substitute rapidly among each other. This trend is evidenced when studying the pattern of PPS substitution in the influenza HA protein. The H1N1 influenza virus entered the human population sometime prior to the 1918 influenza pandemic (21, 22) and has remained largely as a single lineage since then (except from the introduction of a separate lineage of reassortant H1N1 swine virus in 2009 pandemic (23)). Fig. 3 shows the pattern of amino acid substitution for the 18 PPS sites in influenza HA between 1918 and 2009. For example, site 3 remained virtually fixed for alanine between 1918 and the late 1990’s, and then suffered several back and forth substitutions between alanine and valine between the late 1990’s and 2009, while site 142 has been characterised by shifts between lysine and asparagine during that time period. It’s clear from Fig. 3 that the majority of PPS sites in the HA protein are characterised by back-and-forth substitutions among a fairly reduced set of amino acids.

**Fig. 3.**
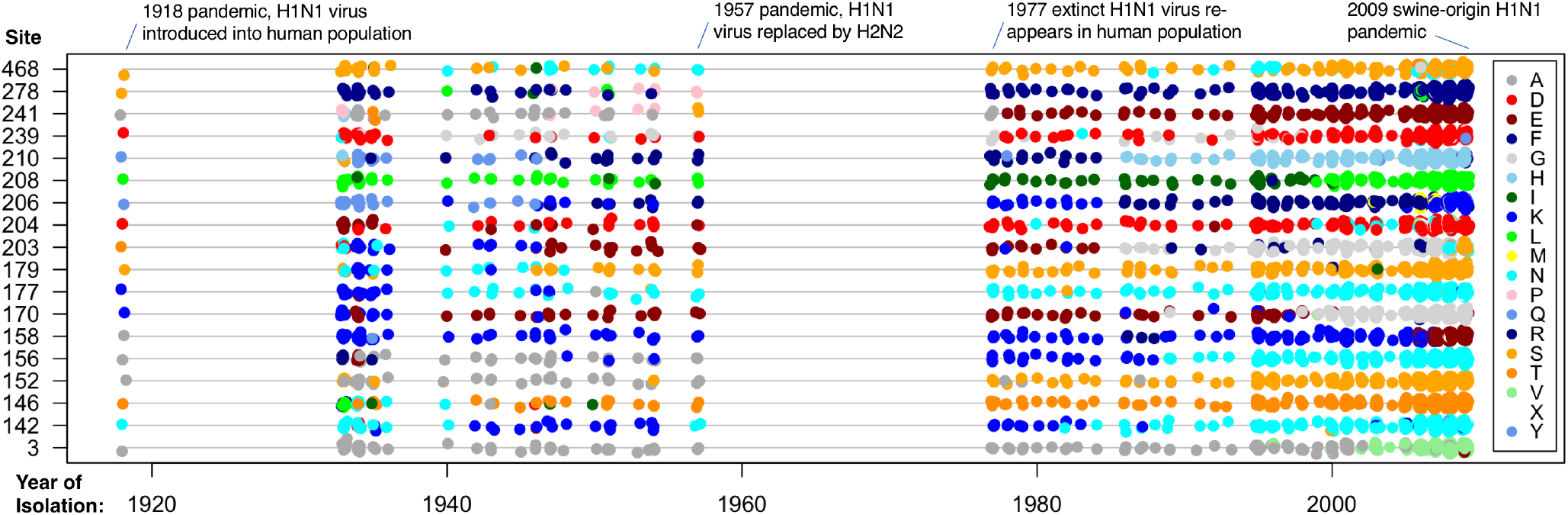
Pattern of amino acid subsitution in PPS sites of human influenza (H1N1) HA protein between 1918 and 2009. Each colored dot represents a particular amino acid as indicated in the legend.

## Discussion

Mutation-selection codon substitution models have been successfully used to study the distribution of selection coefficients in proteins (17, 24), to detect selection shifts during adaptation (25), shifting balance (26), and to understand protein evolution given structural constraints (27), among others. Here we extended the mutation-selection framework to the case of PPS, and we believe the new model can help in gaining insight on the nature of selection on proteins. In particular, we note the PPS model is general and has other models as special cases. For example, when *Z_k_* ≠ 0 and *F_i,j_ = F_j,i_* for all *i, j*, we have

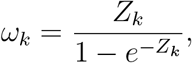

and the model of Eq. 2 can be written as *q_i,j_ = μ_i,j_ω_k_* if the substitution is non-synonymous and *q_i,j_ = μ_i,j_* if otherwise. In other words, the classic codon model of Nielsen and Yang (6, 28) is a special case of Eq. 2 when all codons are assumed to have the same fitness. On the other hand, when *Z_k_* = 0 and *F_i,j_ ≠ F_j,i_*, the model of Eq. 2 reduces to the mutation-selection model of Halpern and Bruno (11).

The PPS model is also flexible, as it appears to have performed well for the different modes of selection studied here. For example, RBCL is the major subunit of the RuBisCo enzyme responsible for the fixation of carbon during photosynthesis. The efficiency of RuBisCo is affected by CO2 concentration in the air, air temperature, humidity, and solar radiation among others. Thus, RBCL has been under persistent adaptive pressures during the successful adaptive radiation of angiosperms around the ecoregions of the world. This is akin to the persistent environment change model originally envisaged by Kimura (1, 12). On the other hand, the influenza HA protein is the classic example of positive selection on a pathogen evading its hosts’ herd immunity, and we showed here the PPS model performed well in detecting this mode of adaptation. We believe the new PPS model, together with previous mutation-selection models that relaxed the assumption of constant fitnesses (25, 29), now encompass the major modes of selection in proteins.

Perhaps the most important insight from the application of the PPS model to real data is that the criteria *ω* > 1 to detect positive selection in proteins is conservative. As we show here, sites under PPS are also under strong purifying constraints, and, at equilibrium, produce many deleterious mutations that are removed by selection. Because *ω_k_* is the weighted average over the rate of all possible synonymous substitutions at a site, it follows that *ω_k_* will be reduced if there are many deleterious mutations at the site even if the site is shifting rapidly among a few positively selection amino acids. We believe this insight should be incorporated into the much faster codon substitution models used in phylogenomic analyses, such as the branch-site model (20), to improve power in detecting adaptation in proteins.

## Materials and Methods

### Maximum penalised likelihood estimation and likelihood ratio test of PPS

The swMutSel model (17, 29) is the special case of swMutSel-PPS when *Z_k_* = 0. We use swMutSel as a null model (*H*_0_ : *Z_k_* = 0) and swMutSel-PPS as the alternative model (*H*_1_ : *Z_k_* > 0) in a likelihood-ratio test. The vector of fitnesses at site *k*, **F***_k_* = (*F_i,k_*) and the PPS component, *Z_k_* are estimated by maximising a penalised likelihood. The penalty on **F***_k_* is the Dirichlet-based penalty of (17), while for *Z_k_* we use an exponential penalty 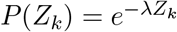, where the regularization parameter, *λ*, controls the strength of the penalty. When *λ* = 0 there is no penalty while *λ* > 0 leads to increasingly stronger penalties on the estimation of *Z_k_*. To speed up computation, the mutational parameters, required to construct *μ_ij_*, and the branch lengths on the phylogeny are estimated under the FMutSel0 model (15) as explained in (17). We recommend the optimisation routine is repeated three times using different parameter start values to ensure convergence to the correct MPLEs. Let the maximum penalised log-likelihood for site *k* be 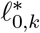 and 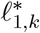, under the *H*_0_ and *H*_1_ hypotheses respectively. The test statistic is the difference in log-likelihoods 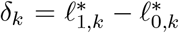. If the test statistic is significantly different from zero, this is evidence site *k* is evolving under PPS. The distribution of the 2*δ_k_* statistic, when the null hypothesis is true, does not follow a *χ*^2^ distribution. There are two reasons for this. First, because *Z_k_* = 0 is at the boundary of parameter space, the test statistic would be, asympototically, distributed as a 50:50 mixture of a *χ*^2^ distribution and a 0.5 point probability mass at 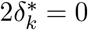 ((30, 31)). The second reason is that the penalty on *Z_k_* affects the 50:50 proportion because the penalty forces the estimates of *Z_k_* towards zero.

Because we do not know what the asymptotic distribution of 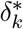 should be, we use Cox’s (18, 19) Monte Carlo simulation to obtain the null distribution of *δ_k_*. For a given site *k* in the alignment, we simulate *N* replicate sites on the phylogeny using the maximum penalised likelihood estimates (MPLEs) of **F***_k_* under *H*_0_ (that is, when *Z_k_* = 0 for all *k*). The distribution, Δ*_k_*, is determined by the difference in log-likelihood between the two models for each simulated site: 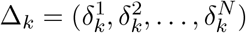 where 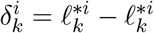 is the log-likelihood difference for the *i*-th simulation. If the test statistic from the real data (*δ_k_*) is larger than, say, 95% of ∆*_k_*, we reject the null hypothesis *H*_0_ (no PPS) and accept the alternative hypothesis *H*_1_ (PPS) at the *α* = 0.05 significance level. When analysing an ensemble of sites in a multiple sequence alignment, we correct for multiple testing using a false discovery rate procedure to select candidate PPS sites (32).

### Pade approximation to calculate the matrix exponential

Calculation of the likelihood along a branch of length *t* in the phylogeny requires calculation of **P**(*t*) = exp *t***Q**_k_, where **Q***_k_* = (*q_ij,k_*) is the substitution matrix (Eq. 2). However, because the PPS model is irreversible, the usual Eigen decomposition algorithm used to calculate **P**(*t*) is not stable (33). Here we use the Pade approximation (34)

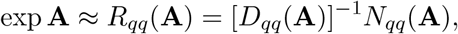

where 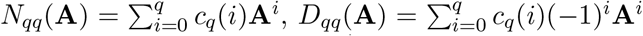, and *c_q_* (*i*) = (2*q − i*)!*q*!*/*(2*q*)!*i*!(*q − i*)!. Note exp **A** = (exp **A***/m*)^*m*^, whith *m* = 2*^j^* for some integer *j*. Accuracy is improved considerably by choosing a suitable *j* such that the Padé approximation works well for exp **A***/m*. Then

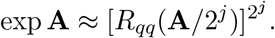

Appropriate values for *q* and *j* are chosen according to the size of **A** and the desired accuracy in the calculation of exp **A** (34).

In our model, the calculation of the likelihood at a site involves multiple computations of exp **Q***t* for every branch in the phylogeny. We choose *q* and *j* according to the largest branch length *t*. Because (*t***Q***/m*)^*i*^ = (*t/m*)^*i*^**Q**^*i*^, we calculate all necessary *c_q_*(*i*) and **Q**^*i*^ once and cache these in memory throughout the likelihood calculation. Calculating **Q**^*i*^ once is more efficient than setting **A = Q***t* and applying the Pade approximation directly. Instead, we compute **B**^*i*^ = (*t/m*)^*i*^**Q**^*i*^, *R_qq_*(**B**), and finally 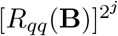 for each value of *t*. We found this matrix exponentiation algorithm is approximately 1.5 times faster than the Taylor series approximation suggested in phylogenetics (33), albeit using more memory to store the precalculated matrix powers.

### Simulated data

To test the specificity and sensitivity of the LRT for PPS, we simulated sites on a balanced 512-taxa tree with branch lengths equal to 0.0125 neutral substitutions per site (17). We simulated sites under a null model with no PPS (*H*_0_ : *Z* = 0), and under the alternative model with PPS (*H*_1_ : *Z* > 0) with three strengths of selection *Z* = 2, 5, 10. Amino acid fitnesses for each site were sampled from a bimodal normal distribution with ten randomly-selected amino acids chosen to have *F ~ N* (0, 1) and the remaining amino acids to have *F ~ N* (−10, 1). The branch lengths and mutation parameters were fixed to their true values (*k* = 2, *π*\* = 0.25) throughout the analysis and only the sitewise fitnesses (**F**) and diversifying selection (*Z*) parameters were estimated. For each simulation setup, we simulated 1,000 sites. Then for each simulation setup we calculated the MPLE and the LRT as described above, using *N* = 100 replicates in Cox’s procedure. In all analyses the Dirichlet penalty on *F_k_* has *α* = 0.01, and three strengths of penalty on *Z* were tested, *λ =*{0.01, 0.5, 1.0}.

Using the LRT results, we determined the false positive and false negative rates. The false positive rate is calculated by determining the number of tests that incorrectly rejected the null hypothesis (*Z* = 0). The true positive rate is calculated from the number of tests that correctly rejected the null hypothesis (*Z* > 0). The results show that the FPR of the LRT is only slightly more than expected at significance level of 0.05 (Table 1). The test has excellent power when strength of selection is moderate or strong (*Z* = 5 or = 10) and is robust to the penalty on *Z* (Table 1).

### Real sequence data

We downloaded 3,120 HA protein-coding sequences of human influenza H1 viruses (excluding 2009 pandemic-H1N1 and partial sequences) from the NIAID Influenza Research Database (35); we downloaded 3,490 RuBisCO eudicotyledon sequences from a previous study (36); and we downloaded CYTB genes of placental mammals from NCBI RefSeq (37) mitochondria genomes. We reduced the HA and RuBisCO datasets to 466 and 478 sequences respectively by using CD-HIT (38) with clustering thresholds of 99.3% and 96% of amino acid sequence identity. The CYTB data was reduced to 418 sequences by keeping one sequence per mammal genus. Sequences were aligned using PRANK (39), and the alignments used to estimate tree topologies with RAxML under the GTRCAT model (40). Because the swMutSel-PPS model is irreversible, trees must be rooted. Thus, outgroups were used to root the trees: Avian influenza (HA), monocotyledons (RuBisCO) and monotremes (CYTB). Outgroups were removed and analyses carried out under on the rooted ingroup tree (for the PPS model), and the unrooted ingroup tree (for the no PPS model). Sites conserved for a single amino acid or with residues in fewer than 50 taxa were not analysed. This corresponds to 31, 23 and 27 sites in the RBCL, HA and CYTB alignments respectively. MPLE and LRT were carried out as described above using *α* = 0.01 and *λ* = 0.001.

### Software Implementation

The mutation-selection PPS codon substitution model is implemented in the swMutSel computer program available at github.com/tamuri.

## ACKNOWLEDGMENTS

MdR is supported by Biotechnology and Biological Sciences Research Council (BBSRC, UK) award BB/T01282X/1.

## Notes

The authors declare no conflict of interest.

### Competing Interest Statement

The authors have declared no competing interest.

